# The usefulness of Nanopore Sequencing in Whole-Genome Sequencing-Based Genotyping of *Listeria monocytogenes* and *Salmonella enterica* serovar Enteritidis

**DOI:** 10.1101/2024.01.29.577746

**Authors:** Yu-Ping Hong, Bo-Han Chen, You-Wun Wang, Ru-Hsiou Teng, Hsiao Lun Wei, Chien-Shun Chiou

## Abstract

Bacterial genotyping through whole-genome sequencing plays a crucial role in disease surveillance and outbreak investigations in public health laboratories. This study assessed the effectiveness of Nanopore Oxford Technologies (ONT) sequencing in the genotyping of *Listeria monocytogenes* and *Salmonella enterica* serovar Enteritidis. Our results indicated that ONT sequences, generated with the R10.4.1 flow cell and basecalled using the Dorado 0.5.0 Super Accurate 4.3 model, exhibited comparable accuracy to Illumina sequences, effectively discriminating among bacterial strains from outbreaks. These findings suggest that ONT sequencing has the potential to be a promising tool for rapid whole-genome sequencing of bacterial pathogens in public health laboratories for epidemiological investigations.

## INTRODUCTION

The advance of next-generation sequencing (NGS) techniques has made whole-genome sequencing (WGS) of bacterial pathogens economically accessible. The WGS approach provides rich data for identifying bacterial serotypes, antimicrobial resistance determinants, virulence genes, and genotyping bacterial strains in epidemiological investigations (1-3). Among the NGS platforms, Illumina sequencing generates large-scale highly accurate sequences, leading to its widespread use in WGS-based genotyping of bacterial pathogens. Nevertheless, Illumina sequencing generates relatively short sequence reads, posing challenges in assembling genomes with repetitive regions and structural variations (4). Furthermore, the long turnaround time and high equipment cost associated with Illumina sequencing impose practical constraints in routine genotyping of bacterial isolates for real-time disease surveillance.

In contrast, Oxford Nanopore Technologies (ONT) sequencing emerges as an alternative with distinctive advantages. ONT sequencing has a rapid turnaround time and lower equipment costs. Moreover, it produces long sequence reads, making it a promising tool for WGS of bacterial isolates. ONT sequencing has proved effective in rapid species identification and detection of virulence and antimicrobial resistance genes (5, 6). Nevertheless, ONT sequencing exhibits lower base accuracy compared to Illumina sequencing, especially in homopolymeric regions, limiting its utility in molecular subtyping of bacterial isolates for disease surveillance and outbreak investigation, where sequence accuracy is exceptionally desired.

Studies have suggested that base modifications can contribute significantly to base calling errors in ONT sequencing (7, 8). Recent developments by ONT, including the introduction of Flowcells (R10.4.1) and Chemistry (SQK-NBD114.24), have demonstrated raw read accuracy exceeding 99.1% (9). Despite this progress, challenges in prokaryotic organism sequencing for bacterial genotyping were noted in a study by Lohde et al. (7). In the study, the researchers indicated that the accuracy of sequences generated using ONT R10.4.1 flow cells and refined with tools available at the time is inadequate for bacteria isolate genotyping in outbreak tracing. Nevertheless, our current study illustrates that ONT sequencing, employing R10.4.1 flow cells and the Dorado 0.5.0 Super Accurate (SUP) 4.3 model, yields sequences with accuracy comparable to Illumina sequencing in the cgMLST and wgSNP analysis of *Listeria monocytogenes* isolates and *Salmonella enterica* serovar Enteritidis isolates from outbreaks.

## MATERIALS AND METHODS

### Bacterial isolates

Twelve *L. monocytogenes* and 23 *S*. Enteritidis isolates were included in this study. *L. monocytogenes* isolates were recovered from sporadic listeriosis cases in hospitals in Taiwan between 2019 and 2020. The *L. monocytogenes* isolates belonged to 8 sequence types, including ST1 ST5, ST87, ST101, ST155, ST378, ST1081, and ST1532 (see Supplemental Material Table S1). Our previous study indicates that 5 of the 12 isolates have numerous base modification-mediated errors in the sequences generated using ONT R9.4 flow cells and the Rapid Barcoding Kit (SQK-RAD004) (8). The *S*. Enteritidis isolates were recovered from 6 foodborne disease outbreaks in the laboratories of the Taiwan Centers for Disease Control (Taiwan CDC) between 2014 and 2022 and genotyped using the standardized PulseNet PFGE protocol (Table S2) (10). The collection of these bacterial isolates was executed through a series of disease surveillance projects, all of which obtained ethical approval from the Institutional Review Board of the Taiwan CDC, Ministry of Health and Welfare. These projects were registered under the IRB Numbers 110109 and 110111.

### Genomic DNA extraction

DNA of bacterial isolates was extracted for WGS using the Qiagen DNeasy blood and tissue kit (Qiagen Co., Germany), manipulated according to the protocol provided by the manufacturer.

### Illumina sequencing

Illumina sequencing was conducted in the Central Region laboratory of Taiwan CDC using the Illumina MiSeq sequencing platform (Illumina Co., USA). Illumina DNA libraries were constructed using the Illumina DNA Prep, (M) Tagmentation Kit (Illumina Co.), and sequencing was run with the MiSeq Reagent Kit v3 (2 × 300 bp), manipulated according to the manufacturer’s instructions. Sequence reads were trimmed using fastp v0.23.0 (11), *de novo* assembled using SPAdes v3.15.3 (12), and polished using POLCA (13).

### ONT sequencing and basecalling

ONT sequencing was performed on the MinION Mk1b (Oxford Nanopore Technologies plc, UK) in the Central Region laboratory. Nanopore DNA libraries for the R9.4 flow cells (FLO-MIN106D) were constructed using the Ligation Sequencing Kit (SQK-LSK109) and the Native Barcoding Expansion Kit (EXP-NBD-114). For the R10.4.1 flow cells (FLO-MIN114), the Rapid Barcoding Kit (SQK-RBK114.24 kit) was used to construct libraries for the rapid sequencing method, and the Native Barcoding Kit (SQK-NBD114.24) was used to construct libraries for the ligation-duplex method. Nanopore raw signal data were basecalled using Dorado 0.5.0 dna_r9.4.1_e8_sup@v3.3, dna_r10.4.1_e8.2_400bps_sup@v4.2.0, and dna_r10.4.1_e8.2_400bps_sup@v4.3.0 models, with or without modification-mediated error correction using the Modpolish toolkit (8).

### Workflow for Assembly and Polishing of ONT sequences

The basecalled ONT reads were trimmed using Porechop v0.2.4 (https://github.com/rrwick/Porechop) to remove adapters, followed by using nanoq v0.10.0 (14) and Filtlong v0.2.1 (https://github.com/rrwick/Filtlong) to filter lengths greater than 10,000 bp and throw away the worst 10% quality reads for downstream assembly. KMC v3.2.1 (15) was applied to estimate genome size and Rasusa v0.7.1 (16) was used to randomly subsample 100x coverage of sequence reads for assembly. Subsampled reads were subjected to *de novo* assembly using Flye v2.9.2 (17), and assembled circular sequences were reoriented using Dnaapler v0.4.0 (18), and polished with medaka v1.11.3 (https://github.com/nanoporetech/medaka). Besides, Plassembler v1.5.0 (19) was used to assemble plasmids.

### cgMLST and wgSNP analysis

Assembled Illumina contigs, Nanopore contigs, and Modpolish-corrected Nanopore contigs were subjected to generate core genome multilocus sequence typing (cgMLST) and whole genome single nucleotide polymorphism (wgSNP) profiles. cgMLST profiling was performed using an in-house-developed cgMLST profiling tool and the cgMLST schemes for *L. monocytogenes* (2,172 core genes) and *Salmonella* (3,241 core genes), available on the openCDCTW/Benga Github repo (https://github.com/openCDCTW/Benga). SNP calling was performed using the Split Kmer Analysis (SKA) toolkit v0.3.5 (20).

### Phylogenetic tree

Phylogenetic trees were constructed with cgMLST or wgSNP profiles using the single linkage algorithm. The correlation between two trees was measured by Baker’s Gamma index (BGI) (21).

### Data availability

The Illumina short sequence reads and ONT R10.4.1 sequence reads through basecalling with the Dorado v0.5.0 SUP Model 4.3 for 12 *L. monocytogenes* and 23 *S*. Enteritidis isolates were deposited under BioPorject accession no. PRJNA428278 in the National Center for Biotechnology Information. The accession numbers for each isolate are listed in the Supplemental Material Table S1 and Table S2.

## RESULTS

### WGS analysis of *L. monocytogenes*

WGS of *L. monocytogenes* isolates were conducted using the Illumina MiSeq, ONT R9.4 with the ligation method, and ONT R10.4 with the ligation-duplex method. Dorado 0.5.0 SUP3.3, 4.2, and 4.3 models were employed for basecalling of ONT reads with or without modification-mediated error correction using the Modpolish toolkit. Significant differences were observed between ONT R9.4 and Illumina sequences, showing an average of 554 (14-1,475) mismatches in cgMLST profiles and 1,496 (0-5,430) SNPs in wgSNP profiles (Table 1). Five isolates (R20.0026, R20.0030, R20.0127, R20.0148, and R20.0150) exhibited particularly high numbers of mismatches. The application of Modpolish resulted in a substantial improvement in ONT R9.4 sequences, reducing mismatches from 1,214-1,475 loci to 1-15 loci in cgMLST profiles for these isolates.

**TABLE 1.**
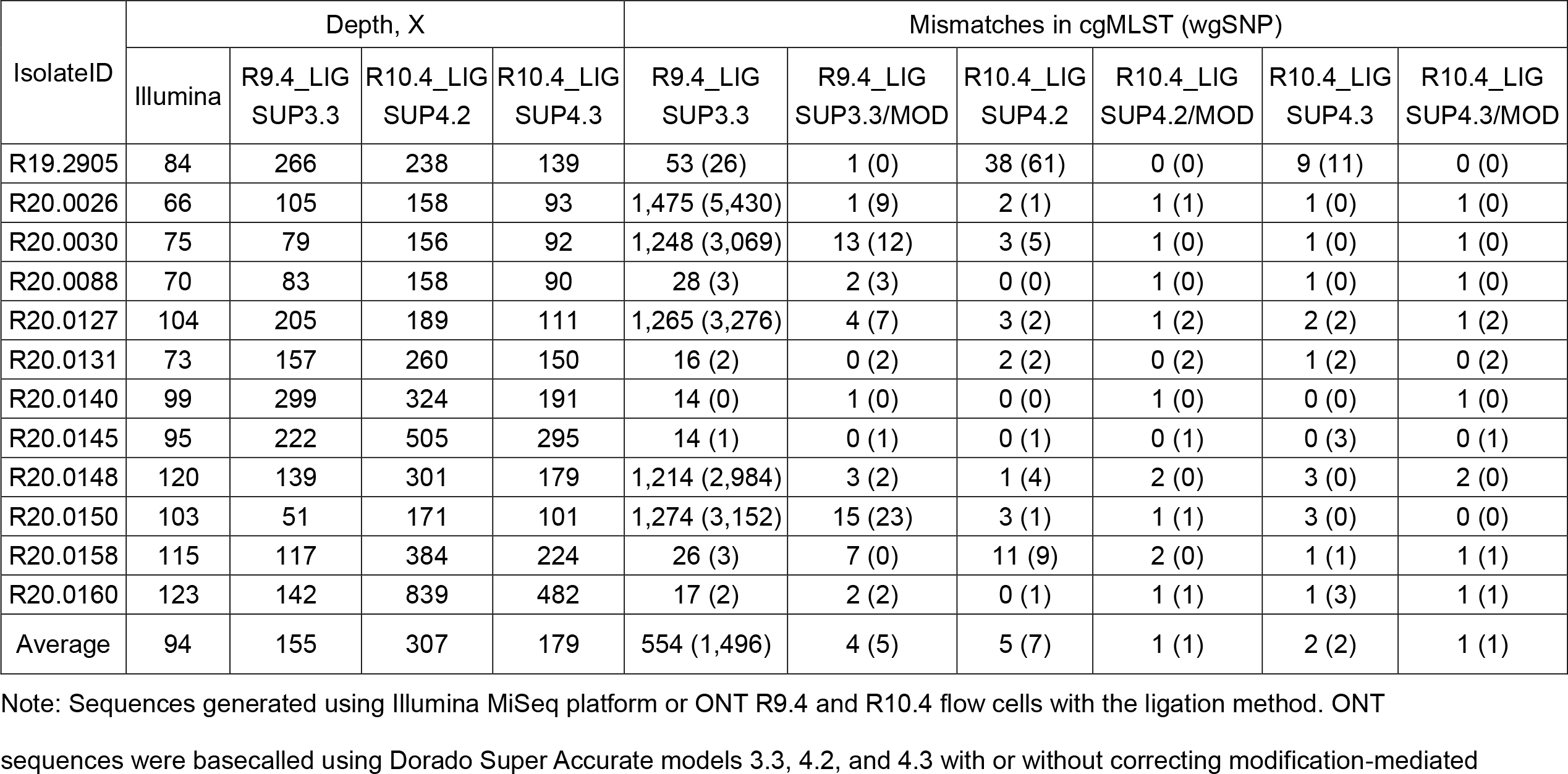
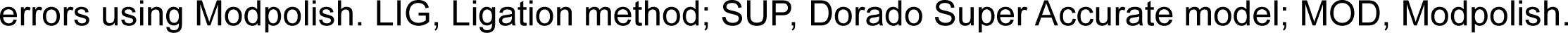
Comparison of cgMLST and wgSNP profiles of *Listeria monocytogenes* isolates generated from Illiumina and ONT sequences

| IsolateID | Depth, X |  |  |  | Mismatches in cgMLST (wgSNP) |  |  |  |  |  |
| --- | --- | --- | --- | --- | --- | --- | --- | --- | --- | --- |
|  | Illumina | R9.4_LIG<br>SUP3.3 | R10.4_LIG<br>SUP4.2 | R10.4_LIG<br>SUP4.3 | R9.4_LIG<br>SUP3.3 | R9.4_LIG<br>SUP3.3/MOD | R10.4_LIG<br>SUP4.2 | R10.4_LIG<br>SUP4.2/MOD | R10.4_LIG<br>SUP4.3 | R10.4_LIG<br>SUP4.3/MOD |
| R19.2905 | 84 | 266 | 238 | 139 | 53 (26) | 1 (0) | 38 (61) | 0 (0) | 9 (11) | 0 (0) |
| R20.0026 | 66 | 105 | 158 | 93 | 1,475 (5,430) | 1 (9) | 2 (1) | 1 (1) | 1 (0) | 1 (0) |
| R20.0030 | 75 | 79 | 156 | 92 | 1,248 (3,069) | 13 (12) | 3 (5) | 1 (0) | 1 (0) | 1 (0) |
| R20.0088 | 70 | 83 | 158 | 90 | 28 (3) | 2 (3) | 0 (0) | 1 (0) | 1 (0) | 1 (0) |
| R20.0127 | 104 | 205 | 189 | 111 | 1,265 (3,276) | 4 (7) | 3 (2) | 1 (2) | 2 (2) | 1 (2) |
| R20.0131 | 73 | 157 | 260 | 150 | 16 (2) | 0 (2) | 2 (2) | 0 (2) | 1 (2) | 0 (2) |
| R20.0140 | 99 | 299 | 324 | 191 | 14 (0) | 1 (0) | 0 (0) | 1 (0) | 0 (0) | 1 (0) |
| R20.0145 | 95 | 222 | 505 | 295 | 14 (1) | 0 (1) | 0 (1) | 0 (1) | 0 (3) | 0 (1) |
| R20.0148 | 120 | 139 | 301 | 179 | 1,214 (2,984) | 3 (2) | 1 (4) | 2 (0) | 3 (0) | 2 (0) |
| R20.0150 | 103 | 51 | 171 | 101 | 1,274 (3,152) | 15 (23) | 3 (1) | 1 (1) | 3 (0) | 0 (0) |
| R20.0158 | 115 | 117 | 384 | 224 | 26 (3) | 7 (0) | 11 (9) | 2 (0) | 1 (1) | 1 (1) |
| R20.0160 | 123 | 142 | 839 | 482 | 17 (2) | 2 (2) | 0 (1) | 1 (1) | 1 (3) | 1 (1) |
| Average | 94 | 155 | 307 | 179 | 554 (1,496) | 4 (5) | 5 (7) | 1 (1) | 2 (2) | 1 (1) |

Note: Sequences generated using Illumina MiSeq platform or ONT R9.4 and R10.4 flow cells with the ligation method. ONT sequences were basecalled using Dorado Super Accurate models 3.3, 4.2, and 4.3 with or without correcting modification-mediated
errors using Modpolish. LIG, Ligation method; SUP, Dorado Super Accurate model; MOD, Modpolish.

The accuracy of ONT sequences was notably enhanced with the use of R10.4 flow cells. ONT R10.4 sequences basecalled using Dorado SUP4.2 exhibited 5 (0-38) mismatches in cgMLST profiles and 7 (0-61) SNPs in wgSNP profiles compared to Illumina sequences. Sequences basecalled with SUP4.3 differed by 2 (0-9) loci in cgMLST profiles and 2 (0-11) SNPs in wgSNP profiles. Further improvement in ONT R10.4 sequence accuracy was achieved through Modpolish. Notably, the differences in cgMLST profiles for R19.2905 and R20.0158 reduced from 38 and 11 loci to 0 and 2 loci, respectively, for ONT R10.4 sequences converted using SUP4.2 and refined with Modpolish (Table 1).

In comparison to ONT R9.4 sequences, ONT R10.4 sequences, when converted using SUP4.2 and SUP4.3 and refined with Modpolish, exhibited higher Qscores, fewer indels, and fewer mismatches (see Supplemental Material Table S1). Regardless of whether basecalling was performed with SUP4.2 or SUP4.3, ONT R10.4 sequencing effectively corrected the high numbers of modification-mediated errors in the sequences generated from ONT R9.4 for the 5 isolates (R20.0026, R20.0030, R20.0127, R20.0148, and R20.0150) (Table 1).

#### WGS analysis of *S*. Enteritidis

WGS analysis was conducted on 23 *S*. Enteritidis isolates from 6 outbreaks using ONT R10.4 flow cells with the Rapid Barcoding and Ligation-duplex (Native Barcoding) kits. Basecalling was performed using Dorado 0.5.0 SUP4.2 and SUP4.3 models, with or without modification-mediated error correction using Modpolish.

Comparisons between ONT sequences generated using the Rapid Barcoding kit with the SUP4.2 model (R10.4_RAP SUP4.2) and Illumina sequences exhibited differences of an average of 61 (20‒160) loci in the cgMLST profiles and 28 (5‒69) SNPs in wgSNP profiles (Table 2). Similarly, ONT sequences generated using the Ligation kit and the SUP4.2 model (R10.4_LIG SUP4.2) exhibited differences of 58 (31‒ 146) loci and 24 (11‒62) SNPs compared to Illumina sequences. However, employing the SUP4.3 model significantly improved the accuracy of both ONT R10.4_RAP and ONT R10.4_LIG sequences. ONT R10.4_RAP SUP4.3 sequences differed from Illumina sequences by 7 (5‒10) loci in cgMLST profiles and 2 (0‒6) SNPs in wgMLST profiles. Similarly, ONT R10.4_LIG SUP4.3 sequences displayed comparable results, with differences of 7 (5‒10) loci in cgMLST profiles and 1 (0‒8) SNPs in wgMLST profiles. However, Modpolish polishing of ONT R10.4_RAP SUP4.3 and ONT R10.4_LIG SUP4.3 sequences did not further enhance their alignment with Illumina sequences. In contrast, Modpolish did improve the concordance between Illumina sequences and both ONT R10.4_RAP and ONT R10.4_LIG sequences basecalled with SUP4.2 (see Supplemental Material Table S2).

**TABLE 2.** Comparison of cgMLST and wgSNP profiles of *Salmonella enterica* serovar Enteritidis isolates generated from Illiumina and ONT sequences

| IsolateID | Depth, X |  |  | Mismatches of alleles in cgMLST (wgSNP) profiles |  |  |  |  |  |
| --- | --- | --- | --- | --- | --- | --- | --- | --- | --- |
|  | Illumina | R10.4_RAP<br>SUP4.3 | R10.4_LIG<br>SUP4.3 | R10.4_RAP<br>SUP4.2 | R10.4_LIG<br>SUP4.2 | R10.4_RAP<br>SUP4.3 | R10.4_LIG<br>SUP4.3 | R10.4_RAP<br>SUP4.3/MOD | R10.4_LIG<br>SUP4.3/MOD |
| C14.0554 | 95 | 317 | 212 | 55 (49) | 70 (55) | 6 (2) | 7 (3) | 8 (2) | 12 (6) |
| C14.0556 | 92 | 250 | 158 | 65 (52) | 81 (62) | 9 (5) | 7 (8) | 15 (13) | 14 (12) |
| C14.0594 | 88 | 295 | 228 | 55 (50) | 87 (60) | 8 (3) | 10 (5) | 10 (5) | 11 (6) |
| CD14.112 | 87 | 402 | 206 | 22 (69) | 31 (16) | 8 (2) | 7 (0) | 9 (4) | 11 (6) |
| CN14.029 | 91 | 294 | 208 | 22 (5) | 38 (11) | 6 (1) | 5 (1) | 8 (6) | 12 (10) |
| NC14.007 | 89 | 254 | 190 | 25 (7) | 38 (16) | 5 (0) | 6 (0) | 13 (10) | 8 (3) |
| R16.5198 | 101 | 76 | 152 | 146 (65) | 52 (18) | 7 (0) | 7 (0) | 13 (8) | 9 (5) |
| R16.5203 | 91 | 166 | 219 | 52 (13) | 42 (12) | 6 (0) | 6 (0) | 10 (8) | 6 (0) |
| R16.5222 | 91 | 226 | 132 | 34 (13) | 90 (21) | 6 (1) | 6 (0) | 13 (10) | 10 (8) |
| R16.5225 | 97 | 154 | 166 | 40 (23) | 58 (19) | 6 (1) | 6 (0) | 6 (1) | 10 (5) |
| R17.0835 | 91 | 257 | 147 | 28 (14) | 146 (49) | 6 (0) | 7 (1) | 6 (3) | 11 (9) |
| R17.0840 | 96 | 59 | 139 | 97 (30) | 91 (28) | 8 (1) | 10 (1) | 11 (9) | 8 (6) |
| R17.0847 | 97 | 327 | 196 | 23 (8) | 46 (26) | 8 (5) | 6 (6) | 8 (8) | 8 (6) |
| R19.1081 | 83 | 65 | 149 | 114 (43) | 37 (14) | 8 (6) | 5 (0) | 9 (8) | 6 (3) |
| R19.1090 | 94 | 198 | 125 | 30 (9) | 72 (18) | 6 (1) | 6 (1) | 7 (4) | 8 (8) |
| R19.1092 | 109 | 239 | 172 | 20 (6) | 45 (19) | 6 (0) | 7 (0) | 6 (0) | 6 (0) |
| R19.1218 | 80 | 140 | 201 | 46 (20) | 37 (15) | 6 (1) | 7 (1) | 9 (8) | 6 (3) |
| R19.1220 | 76 | 126 | 223 | 82 (14) | 63 (25) | 6 (1) | 6 (1) | 12 (8) | 6 (3) |
| R19.1468 | 77 | 103 | 206 | 79 (17) | 40 (12) | 10 (1) | 6 (0) | 7 (4) | 5 (3) |
| R19.1471 | 77 | 167 | 213 | 43 (17) | 49 (15) | 6 (2) | 6 (1) | 5 (2) | 6 (1) |
| R22.0060 | 75 | 57 | 206 | 160 (62) | 50 (20) | 5 (2) | 6 (0) | 11 (11) | 6 (0) |
| R22.0063 | 81 | 93 | 178 | 97 (28) | 40 (19) | 7 (1) | 7 (1) | 9 (6) | 7 (2) |
| R22.0075 | 73 | 124 | 162 | 74 (21) | 41 (11) | 6 (1) | 5 (0) | 9 (7) | 7 (1) |
| Average | 88 | 191 | 182 | 61 (28) | 58 (24) | 7 (2) | 7 (1) | 9 (6) | 8 (5) |

Note: Sequences generated using Illumina MiSeq platform or R10.4 flow cells with rapid barcoding or ligation methods and basecalled using Dorado 0.5.0 Super Accurate models 4.2 and 4.3 with or without correcting modification-mediated errors using Modpolish. RAP, rapid barcoding method; LIG, ligation-duplex method; SUP, Dorado 0.5.0 Super Accurate model; MOD, Modpolish.

#### Phylogenetic analysis

Clustering analysis of cgMLST and wgSNP profiles was conducted to assess the similarity between phylogenetic trees constructed with Illumina and ONT sequences, measured by BGI values ranging from -1 to 1. In the case of *L. monocytogenes*, exceptionally high BGI values (0.9999 and 1) were observed for both cgMLST and wgSNP trees constructed using the ONT R10.4 sequences from SUP4.2 and SUP4.3 basecalling, as well as the ONT R10.4 sequences refined with Modpolish (see Supplemental Material Table S3).

For *S*. Enteritidis isolates, the cgMLST trees constructed with ONT R10.4_RAP and ONT R10.4_LIG sequences from SUP4.3 basecalling exhibited significantly higher BGI values compared to sequences from SUP4.2 basecalling (BGI, 0.9983 and 0.9975 vs. 0.3575 and 0.5638) (Supplemental Material Table S3). wgSNP analysis obtained even higher similarity between trees, as indicated by elevated BGI values. Notably, the application of Modpolish enhanced the BGI values for cgMLST and wgSNP trees constructed with ONT sequences from SUP4.2 but not SUP4.3 basecalling.

In intra-outbreak analyses, differences in cgMLST profiles generated from Illumina sequences ranged from 1 to 3 loci among isolates from an outbreak, while those from ONT R10.4_LIG SUP4.3 sequences ranged from 1 to 4 loci (Fig. 1). In wgSNP profiles, Illumina sequences displayed differences of 0 to 4 SNPs among isolates from an outbreak, while ONT sequences displayed differences of 0 to 8 SNPs. Similarly, ONT R10.4_RAP SUP4.3 sequences displayed a high degree of similarity, as evidenced by BGI values of 0.9983 for the cgMLST trees and 0.9984 for the wgSNP trees (see Supplemental Material Table S3).

**FIG 1.**
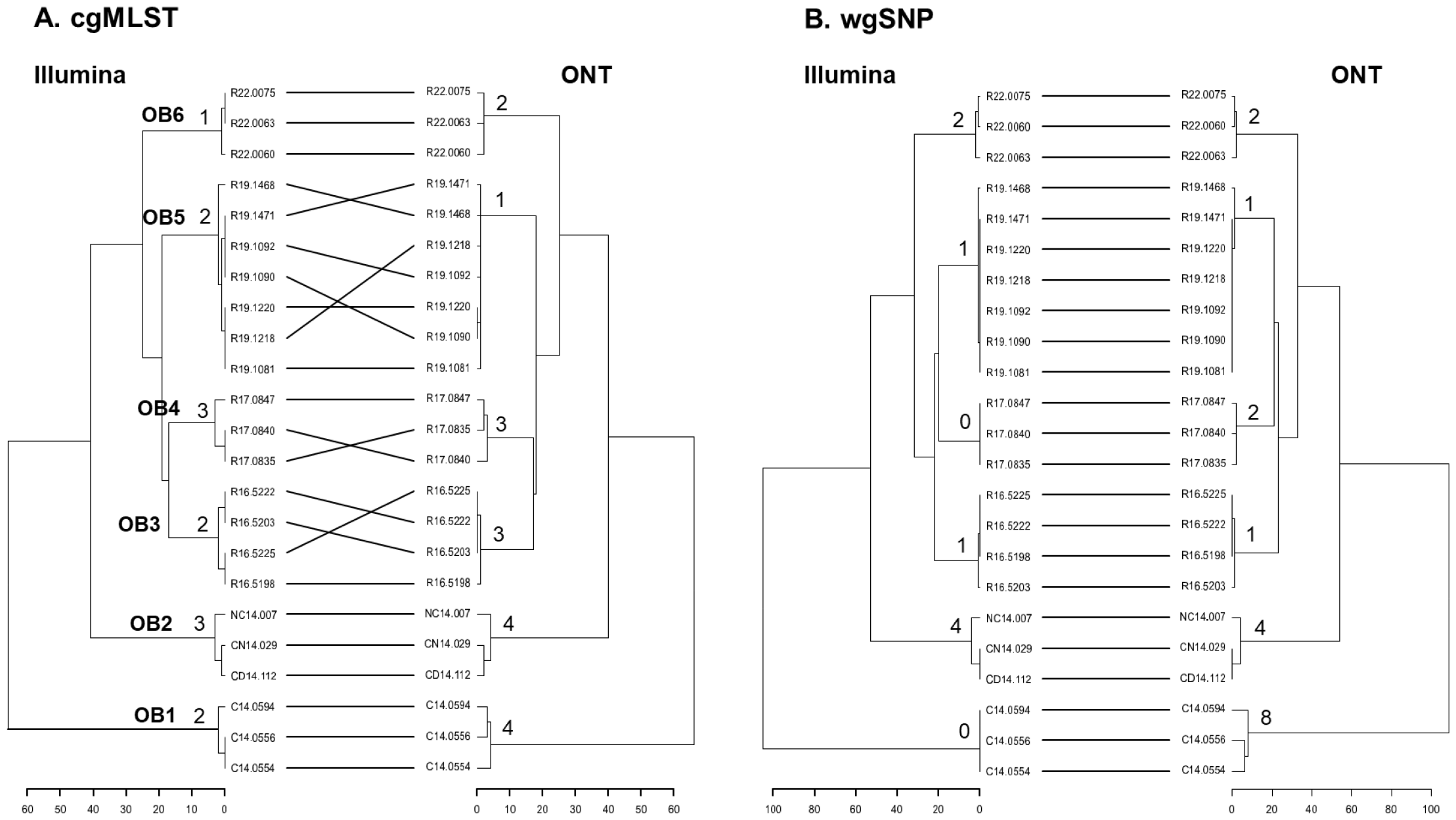
Dendroscope tanglegram comparison between cgMLST trees (A) and wgSNP trees (B), constructed with Illumina contigs and ONT contigs for *Salmonella enterica* serovar Enteritidis isolates from 6 outbreaks.

## DISCUSSION

Our findings indicate that the accuracy of ONT R10.4 sequences from *L. monocytogenes* and *S*. Enteritidis isolates, when subjected to basecalling using the Dorado 0.5.0 Super Accurate model 4.3, closely approximates the accuracy observed in Illumina sequences. Additionally, the accuracy of *L. monocytogenes* ONT sequences can be further improved through the correction of the Modpolish toolkit. Notably, the accuracy of *S*. Enteritidis ONT sequences, generated from both Rapid and Ligation methods, closely parallels that of Illumina sequences. This improvement suggests that ONT sequencing has the potential to be a promising tool for rapid WGS-based genotyping of bacterial strains, thereby greatly contributing to disease surveillance and outbreak investigation practices.In our earlier investigation, we showed that the sequences generated with the ONT R9.4 device lacked the necessary accuracy for WGS-based genotyping of *L. monocytogenes* (8). The primary cause of the inaccuracy was errors resulting from base modification (8). While the Modpolish toolkit effectively proved most of these errors, the polished sequences still failed to meet the required accuracy for WGS-based genotyping (8). In the present study, we demonstrate a notable improvement with the implementation of the ONT R10.4 device along with the Dorado basecaller, effectively eliminating errors attributed to base modifications observed in ONT R9.4 sequences (Table 1). In addition, we demonstrate further improvement in the accuracy of ONT sequences of *L. monocytogenes* through the application of the Modpolish toolkit.

Our data indicate that ONT R10.4 sequences of *S*. Enteritidis, generated through both Rapid and Ligation methods and basecalled using the Dorado SUP4.3 model, exhibit an accuracy comparable to the Illumina sequences. In our initial analysis, we basecalled the ONT R10.4-RAP and ONT R10.4-LIG sequences from *S*. Enteritidis using the Dorado SUP4.2 model, resulting in sequences that did not match the accuracy of Illumina sequences (Tabe 2). Subsequent re-analysis, following the release of SUP4.3, demonstrated a substantial improvement in the accuracy of the ONT R10.4 sequences. Notably, the ONT sequences differ from the Illumina sequences by an average of 7 loci in the cgMLST profiles and 1 to 2 SNPs in the wgSNP profiles (Table 2). The phylogenetic trees constructed for the 23 *S*. Enteritidis isolates using ONT R10.4-RAP and ONT R10.4-LIG sequences closely align with those built using the Illumina sequences (Fig. 1, Supplemental Material Table S3). These findings suggest that ONT sequencing alone may serve as a reliable tool for WGS-based genotyping of bacterial strains in public health laboratories for disease surveillance and outbreak tracing.

We demonstrate the superiority of the Dorado SUP4.3 model over SUP4.2 in converting ONT R10.4 sequences. The refined ONT sequences from both *L. monocytogenes* and *S*. Enteritidis exhibit comparable accuracy to Illumina sequences, as evident in the phylogenetic analysis and outbreak identification (Fig. 1). While our study was conducted only on two bacterial species, a crucial consideration arises regarding the applicability of the SUP4.3. In an investigation, the functionality of the SUP4.2 and SUP4.3 models was assessed using 12 standard genomes representing bacterial species, including *Campylobacter jejuni, C. lari, Escherichia coli, Listeria ivanovii, L. monocytogenes, L. welshimeri, S. enterica, Vibrio cholerae*, and *V. parahaemolyticus* (https://rrwick.github.io/2023/12/18/ont-only-accuracy-update.html). This assessment demonstrates that the SUP4.3 model significantly improved both read accuracy and assembly accuracy compared to the SUP4.2, thereby extending its potential applicability across diverse bacteria species.

In conclusion, ONT sequences generated from R10.4 flow cell and basecalled using the Dorado 0.5.0 SUP4.3 model exhibited accuracy comparable to Illumina sequences in WGS-based genotyping of *L. monocytogenes* and *S*. Enteritidis isolates. However, to comprehensively assess the effectiveness of this method in disease surveillance and outbreak investigation, it is imperative to conduct further investigations involving more isolates from diverse bacterial species.

## ACKNOWLEDGMENTS

This study was funded by the Ministry of Health and Welfare, Taiwan (grant numbers MOHW110-CDC-C-315-113107, MOHW111-CDC-C-315-123108, MOHW112-CDC-C-315-133116, and MOHW113-CDC-C-315-144316).

